# Impaired astrocyte-to-neuron cholesterol trafficking drives synaptic dysfunction in Rett syndrome

**DOI:** 10.64898/2026.07.29.741426

**Authors:** Francesca M. Postogna, Noemi Giancroce, Cecilia Cabasino, Fabio Biella, Ottavia M. Roggero, Martina Breccia, Laura Morelli, Diego Colombo, Alessandro Arcari, Giulia Lunghi, Manuela Valsecchi, Elena Chiricozzi, Nicoletta Landsberger, Marta Valenza, Angelisa Frasca

## Abstract

Rett syndrome (RTT) is a severe X-linked neurodevelopmental disorder caused by loss-of-function mutations in the *MECP2* gene and characterized by profound impairments in neuronal maturation and synaptic connectivity. Increasing evidence indicates that astrocyte dysfunction contributes to RTT pathogenesis through non-cell-autonomous mechanisms, although the molecular pathways underlying defective astrocyte-neuron communication are only partially understood. Astrocytes are the primary source of cholesterol in the brain and support neuronal maturation and synaptic function by supplying cholesterol through ApoE-containing lipoproteins. Although alterations in brain cholesterol metabolism have been reported in RTT, the underlying cellular mechanisms and their functional consequences remain poorly investigated. Here, we studied cholesterol homeostasis in *Mecp2* knock-out (KO) astrocytes and its impact on neuron-astrocyte communication. *Mecp2* KO astrocytes exhibited reduced nuclear localization of the transcriptional regulator Srebp2, together with the downregulation of genes involved in cholesterol biosynthesis and transport. These molecular alterations were associated with intracellular cholesterol and desmosterol accumulation, reduced Abca1 expression and defective ApoE lipidation, despite preserved ApoE expression and cholesterol secretion. Importantly, similar alterations were detected in acutely isolated astrocytes and in the cerebral cortex of *Mecp2* deficient mice, demonstrating that impaired cholesterol homeostasis extends beyond *in vitro* models. Functionally, cholesterol supplementation of astrocyte-conditioned medium rescued the synaptic defects induced in wild-type neurons by *Mecp2* KO astrocytes. Moreover, cholesterol treatment restored pre- and post-synaptic density, as well as axon initial segment length in *Mecp2* heterozygous (HET) neurons. Together, these findings identify defective astrocyte-to-neuron cholesterol trafficking as a key mechanism contributing to neuronal dysfunction in RTT and suggest that strategies aimed at restoring cholesterol functional availability might represent a promising therapeutic avenue for RTT.

## Introduction

Rett syndrome (RTT) is a progressive neurodevelopmental disorder that affects about 1 in 10,000 live female births worldwide, representing the most frequent genetic cause of severe intellectual disability in girls (Chahrour & Zoghbi, 2007; Gold et al., 2024). Loss-of-function mutations in the X-linked *MECP2* gene, encoding the Methyl-CpG-binding protein 2 (MeCP2), are responsible for about 95% of typical RTT cases, and the type of mutation, together with the pattern of X chromosome inactivation and the presence of modifier genes, determine the phenotypic variability among RTT patients (Amir et al., 1999; Chae et al., 2004; Christodoulou & Weaving, 2003). MeCP2 plays crucial roles in CNS development and maintenance and, therefore, its loss of function is associated with severe neurobiological alterations. By analysing *Mecp2* mutant animals and RTT post-mortem tissues, a reduction in brain volume was reported, mainly attributed to defects in neurons (Carli et al., 2023; Chahrour & Zoghbi, 2007; Naidu et al., 2001). Indeed, RTT neurons show smaller soma and reduced dendritic arborisation compared to healthy neurons (Belichenko et al., 2009; Fukuda et al., 2005; Jentarra et al., 2010; Perego et al., 2022). Moreover, the severe and diffuse synaptic alterations, both at morphological and functional levels, have led to considering RTT as a synaptopathy (Boggio et al., 2010; Calfa et al., 2015; Dani et al., 2005; Fukuda et al., 2005). Although molecular changes induced by *MECP2* loss in neurons are considered the primary drivers of synaptic dysfunctions, recent studies have also demonstrated a role for non-cell-autonomous mechanisms largely mediated by astrocytes, which strongly influence neuronal maturation and function (Allen & Eroglu, 2017; Postogna et al., 2025). The involvement of astrocytes in RTT pathogenesis was first reported by *in vitro* studies showing that human and mouse RTT astrocytes failed to support neuronal development, leading to aberrant dendritic morphology (Ballas et al., 2009; Williams et al., 2014). Consistently, selective reactivation of *Mecp2* expression in astrocytes of *Mecp2* KO mice resulted in a significant rescue of both behavioral and neuronal phenotype, highlighting astrocytes as important contributors to disease pathogenesis and potential therapeutic targets (Lioy et al., 2011). To identify the mechanisms underlying astrocyte-mediated neuronal dysfunction, several studies have characterized the molecular and functional alterations associated with *Mecp2* deficiency, reporting defects in multiple processes, including BDNF regulation, cytokine production, cell maturation, morphology and metabolism (Albizzati et al., 2022, 2024; Maezawa et al., 2009; Pacheco et al., 2017; Sun et al., 2023). Among the mechanisms proposed, increasing evidence points to the dysregulation of astrocyte-secreted factors. Indeed, the observation that conditioned medium from *Mecp2* deficient astrocytes is sufficient to impair neuronal development suggests that soluble factors are major mediators of astrocyte-neuron communication in RTT (Albizzati et al., 2024; Ballas et al., 2009). According to this hypothesis, proteomic studies conducted on *Mecp2* KO astrocytes’ secretome identified several dysregulated proteins, including Lcn2, Lgals3 and Igfbp2 (Caldwell et al., 2022; Ehinger et al., 2021). We recently demonstrated that *Mecp2* deficient astrocytes secrete excessive amounts of interleukin-6 (IL-6), leading to reduced synapse density and impaired synaptic function in wild-type (WT) neurons (Albizzati et al., 2024). Nevertheless, although most studies have focused on the increased release of neurotoxic molecules, neuronal dysfunction might also result from insufficient production of astrocyte-derived synaptogenic factors. The most studied synaptogenic factors released by astrocytes include thrombospondins 1 and 2 (TSP1/2), glypican 4 and 6 (GPC4-6), Tumor Necrosis Factor α (TNF-α), Brain-Derived Neurotrophic Factor (BDNF), Hevin, Secreted Protein Acidic and Rich in Cysteine (SPARC), protocadherin and cholesterol (Baldwin & Eroglu, 2017; Chung et al., 2015; Shan et al., 2021). It is well established that in the adult brain, astrocytes represent the main source of cholesterol, which is biosynthesized through the Bloch pathway and exported together with phospholipids by ATP-binding cassette (ABC) transporters, mainly ABCA1 and ABCG1, to lipidate ApoE and generate ApoE-containing lipoproteins that deliver cholesterol to other cells, particularly neurons. Astrocyte-released cholesterol is essential for neuronal development and maturation, as suggested by dendritic spines and synapse degeneration resulting from cholesterol depletion in astrocytes (Goritz et al., 2005; J.-P. Liu et al., 2010; Mauch et al., 2001; Pfrieger, 2003). Multiple lines of evidence suggest that cholesterol metabolism is altered in RTT. The first indication came from a genetic modifier screen in *Mecp2* mutant mice performed by Buchovecky and colleagues. In their search for genes capable of modulating RTT-associated phenotypes, they identified a nonsense mutation in *Sqle*, encoding squalene epoxidase, a key rate-limiting enzyme of the cholesterol biosynthetic pathway. By analyzing symptomatic *Mecp2* null animals, they reported high levels of cholesterol in serum and whole brain homogenates, although the amounts of metabolic intermediates were decreased (Buchovecky et al., 2013a). A subsequent study reported a significant reduction in the rate of cholesterol synthesis in the *Mecp2* null brain starting from P21 and maintained until P56, with a concomitant reduction in cholesterol precursors without observing a reduction in the total amount of cholesterol at any time points (Lopez et al., 2017; Lütjohann et al., 2018). Similarly, studies in RTT patients reported alterations in cholesterol metabolism. Lipidomic analyses performed by liquid chromatography-tandem mass spectrometry (LC-MS/MS) on CSF and plasma of RTT patients revealed reduced cholesterol levels in the CSF, but not in plasma, compared with age-matched healthy controls (Zandl-Lang et al., 2022). Furthermore, a general deregulation of the protein network of cholesterol homeostasis, including HMGCR and SREBP1 and 2 expression, was described in RTT fibroblasts (Segatto et al., 2014).

However, to date no study has investigated whether defects in cholesterol metabolism contribute to impaired astrocyte-neuron crosstalk in RTT or whether cholesterol supplementation could represent a potential therapeutic strategy for this neurodevelopmental disorder.

In this work, we provide the first comprehensive characterization of cholesterol metabolism in *Mecp2* KO astrocytes. We found that *Mecp2* deficient astrocytes exhibit reduced nuclear translocation of the master regulator Srebp2, leading to a general downregulation of genes involved in cholesterol biosynthesis and trafficking. Although cholesterol secretion remained unchanged, *Mecp2* deficient astrocytes accumulated intracellular cholesterol and desmosterol, exhibited reduced Abca1 expression, and released reduced levels of lapidated ApoE, indicating impaired cholesterol trafficking and functional delivery to neurons. Finally, we demonstrate for the first time that cholesterol supplementation improves synaptic morphology in RTT neurons, supporting astrocytic cholesterol metabolism as a potential therapeutic target.

## Methods

### Animal models and genotyping

All animal procedures were performed in accordance with the European Union Communities Council Directive (2010/63/EU) and Italian laws (D.L.26/2014) and approved by the Italian Council on Animal Care (Italian government decree No. 187/2022). Mice were housed in a 12-hours light/12-hours dark cycle with food and water *ad libitum* in an environment maintained at a controlled temperature and humidity. Mecp2*^tm1.1/Bird^* (Gigli et al., 2016) and Mecp2*^Y120D^* (Gandaglia et al., 2019) mouse strains were used. Mice genotyping was obtained through polymerase chain reaction (PCR) on genomic DNA derived from either ear punch, tail or paw biopsies. To extract DNA from samples we used the Phire animal tissue direct PCR kit (#F140WH, ThermoFisher Scientific). PCR was performed on genomic DNA using the primers reported in Table 1.

**Table 1.**
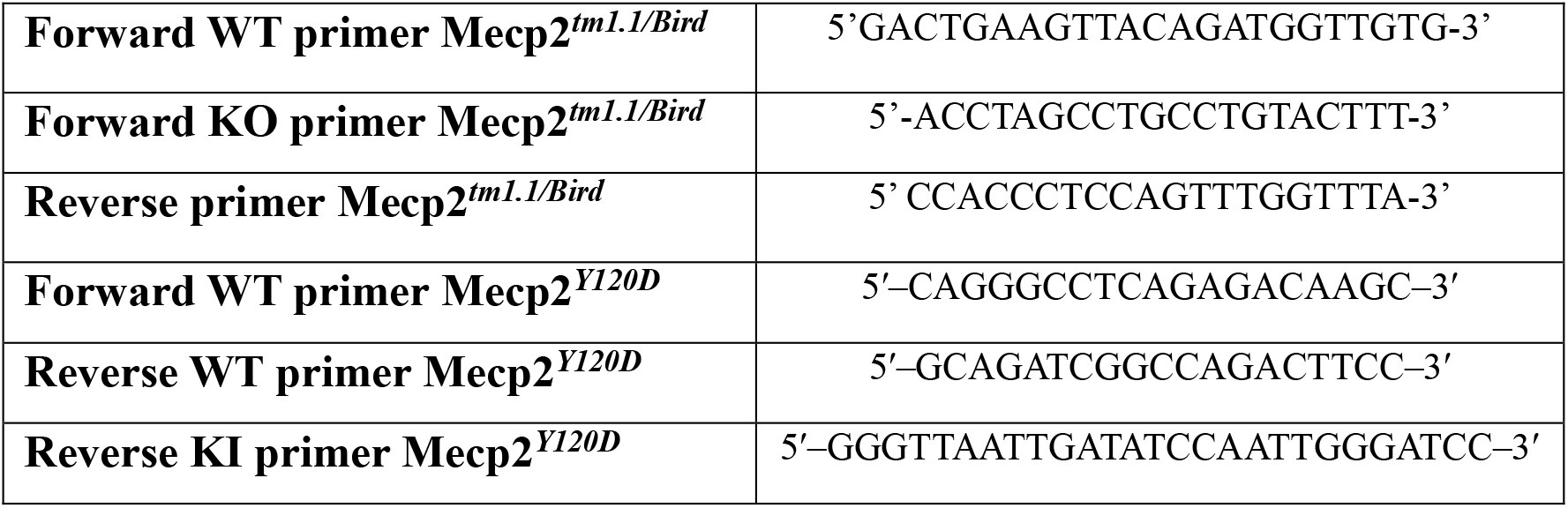
List of primers used for animal genotyping.

### Primary cultures of cortical astrocytes

Primary cultures of astrocytes were obtained from the cortices of P1-P3 (post-natal day) WT and *Mecp2* KO CD1 mice, as previously described (Albizzati et al., 2024). Briefly, cerebral cortices were maintained on ice in Hanks’ Balanced Salt Solution (HBSS; #14175-095, ThermoFisher Scientific), containing 10 mM Hepes, 4 mM Na2HCO3, 1% Penicillin/Streptomycin (P/S; #P0781, Merck) until processing. Tissues were washed with HBSS and then incubated with 0.25% Trypsin/EDTA (#25200-056, ThermoFisher Scientific) for 30 minutes at 37°C. Trypsin was then inactivated by adding astrocytes culture medium, composed by Dulbecco’s modified Eagle medium (DMEM; #41966029, ThermoFisher Scientific), Ham’s F-10 Nutrient mix (#31550023, ThermoFisher Scientific), 10% Fetal Bovine Serum (FBS; # 10500064, ThermoFisher Scientific) and 1% P/S, and tissues were washed three times in culture medium. Samples were dissociated and cell suspension was filtered through a cell strainer of 40 μm pore size. After centrifugation for 6 minutes at 1,500 rpm, cells were resuspended in culture medium and then seeded in a T75 flask, previously coated with 15 ug/ml Poly-D-lysine hydrobromide (PDL; #P7886, Merck). Cells were incubated at 37°C and 5% CO2. At DIV4 (day *in vitro*), flasks were shaken for 6 hours at 37°C in culture medium supplemented with 10 mM Hepes, thus promoting detaching of microglia and oligodendrocytes. Medium was then replaced by fresh culture medium, which was changed every 3 or 4 days. When confluent, cells were washed with Dulbecco’s phosphate-buffered saline (D-PBS; #ECB4004L, Euroclone) and detached by using 0.25% trypsin/EDTA diluted 1:3 in HBSS, for 2 minutes at 37°C. Trypsin was inactivated by subsequent addition of medium, and cell suspension was centrifuged at 1,500 rpm for 6 minutes. Cells were suspended in culture medium and counted in a Bürker chamber with Trypan blue. Then, cells were seeded on a 6-well plate or 100 mm Petri dish coated with PDL (0.1 mg/ml). For immunofluorescence (IF) experiments, astrocytes were seeded on Poly-D-Lysine German Glass Coverslips (*#*1212478, Kleinfeld).

Alternatively, cortical astrocytes were isolated from P7 and P40 *Mecp2* KO and WT mice following the Miltenyi Biotec protocol “Isolation and cultivation of astrocytes from adult mouse brain” (#130107677, Miltenyi Biotec). Cortices from both hemispheres were dissected after careful removal of skin, cartilage and meninges, and tissues were maintained in HBSS until processing. Samples dissociation was obtained through the gentleMACS™ Dissociator with Heaters, which enables mechanical dissociation during the on-instrument enzyme incubation. After dissociation, the Debris Removal Solution and the Red Blood Cell Removal Solution were used to remove myelin, cell debris and erythrocytes. The anti-ACSA-2 Microbead Kit (#130095826, Miltenyi Biotec) was used to isolate astrocytes from the single-cell suspension. 500 μL of PureZOL were added to each sample, which were stored at -80°C until RNA extraction. Alternatively, astrocytes were suspended in pre-warmed AstroMACS Medium (#130117031, Miltenyi Biotec), composed by MACS Neuro Medium, 0.2% AstroMACS Supplement, 2% MACS NeuroBrew-21, 0.25% L-glutamine 0.5 mM (#G7513, Merck) and 1% P/S, and plated on 24-well plate, which was previously coated with 0.1 mg/ml PDL overnight (O/N) at room temperature (RT), followed by 10 µg/ml laminin, for 2 hours at 37°C. Astrocyte medium was half-changed every other day for 3 weeks, corresponding to a DIV when astrocytes reach their classical star-like morphology.

### Primary cultures of cortical neurons

Primary cultures of cortical neurons were generated from WT and *Mecp2* heterozygous (HET) embryos at E15.5 (embryonic day), as already described (Albizzati et al., 2024; Frasca et al., 2020). Briefly, dissected cortices were washed in HBSS before enzymatic dissociation in 0.25% Trypsin/EDTA for 7 minutes at 37°C. Trypsin was then removed by washing each sample three times in HBSS. Mechanical dissociation was performed in DMEM, containing 10% FBS, 1% glutamine and 1% P/S, by gently pipetting. Cells were counted in an automated cell counter (Countess Automated Cell Counter, ThermoFisher Scientific) by using Trypan blue staining, and then cultured in neuron culture medium, composed by Neurobasal plus medium (#A3582901, ThermoFisher Scientific), 2% B27 Plus (#A3653401, ThermoFisher Scientific), 1% P/S. Neurons were plated on PDL glass coverslips and multi-well plates coated with PDL (0.1 mg/ml) and incubated at 37°C and 5% CO2.

### Astrocyte-Conditioned Medium (ACM) preparation

Astrocytes derived from pups were plated at a density of 100,000 cells/well in 6-well plates. When confluent, they were starved by removing serum from the culture medium. In detail, after two washes with DMEM, astrocytes were incubated for 48 hours with neuron culture medium, which was collected, centrifuged for 5 minutes at 160g at 4°C and stored at -80°C until use. ACM samples were used for cholesterol quantification and for treatment of neurons. In this case, ACM was warmed up to 37°C and then added 1:1 to neuronal culture medium for 24 hours (from DIV13 to DIV14). For western blot analyses, ACM was concentrated 10-fold by using Amicon Ultra-4 filter devices (#UFC803008, Merck). ACM was also prepared from MACS-isolated astrocytes. After 3 weeks in culture, the medium was fully replaced with fresh AstroMACS Medium, which was collected after 48 hours, centrifuged for 5 minutes at 1,200 rpm at 4 ° C and used for cholesterol quantification.

### Quantitative Reverse-Transcriptase PCR (qRT-PCR)

Total RNA from murine astrocytes and cerebral tissues was extracted using PureZol (#7326890; Bio-Rad), quantified using a NanoDrop spectrophotometer (ThermoFisher Scientific) and reverse transcribed using the RT2 First Strand Kit (#330404; Qiagen) as instructed by the manufacturer. The resulting cDNA was used as a template for qRT-PCR with SYBR Green Master Mix (#4472908; Applied Biosystems) with designated primers (**Table 2**). Due to the low amount of RNA extracted from MACS-isolated astrocytes, a preamplification step was performed using SsoAdvanced PreAmp Supermix (#1725160, Bio-Rad) prior to qRT-PCR, following the manufacturer’s instructions. Melting curves showed a single product peak, indicating good product specificity. Hprt, Cyclophilin A and/or Rpl13 served as internal standards and fold change in gene expression was calculated using the 2^(-Delta Ct) method.

**Table 2.**
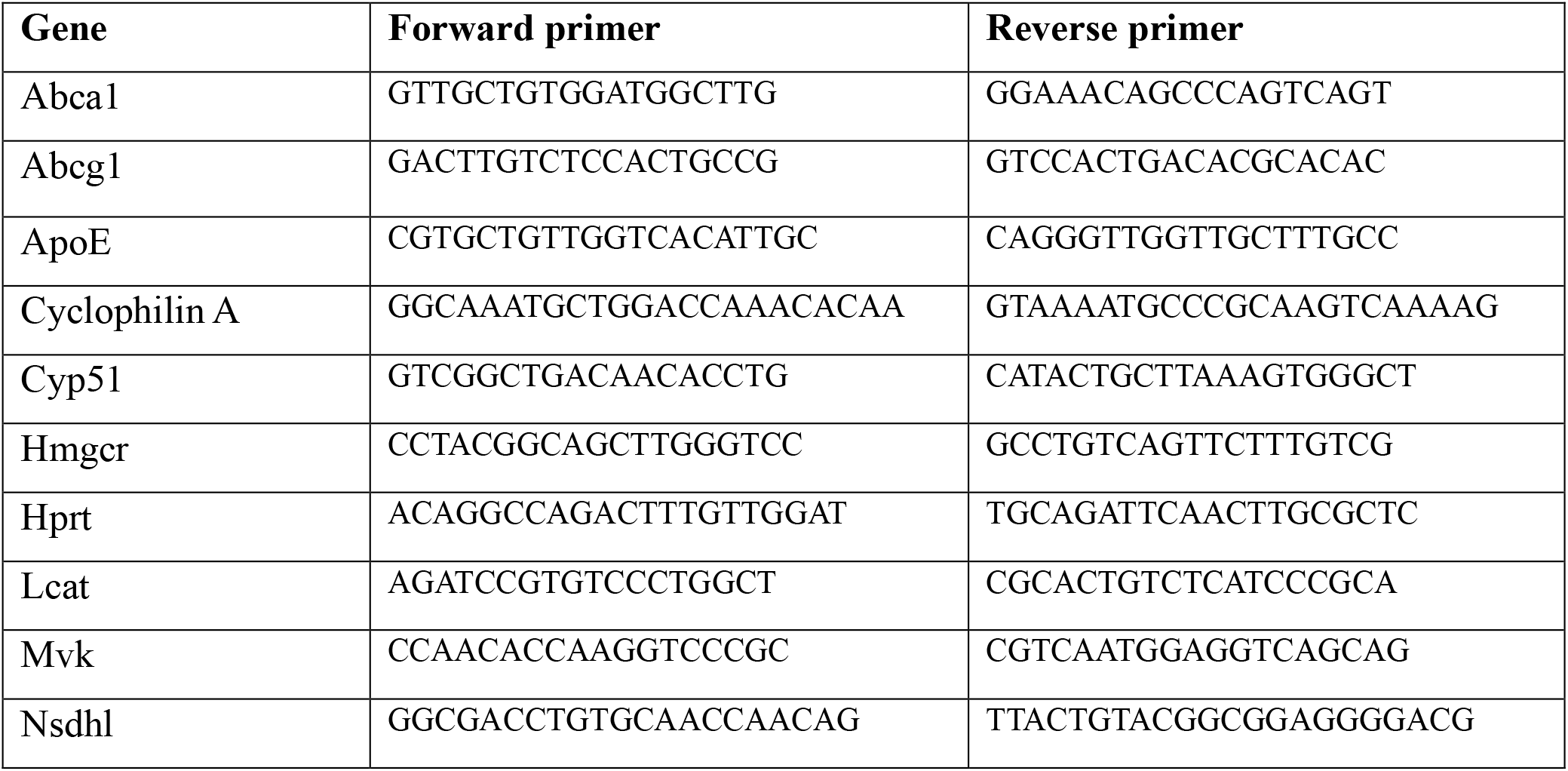
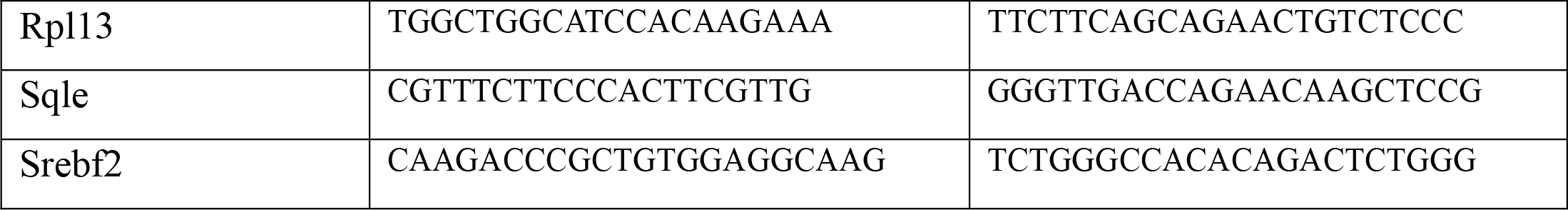
List of the genes tested by qRT-PCR and their respective primers.

### Western blot analysis

Primary astrocyte and cerebral tissues were lysed in ice-cold T-PER^TM^ Tissue Protein Extraction Reagent (#78510, ThermoFisher Scientific) supplemented with protease and phosphatase inhibitor cocktail (#78444, ThermoFisher Scientific). The homogenates were centrifuged for 30 minutes at 160g at 4°C and protein concentrations were measured in the supernatants by BCA assay (#23228, ThermoFisher Scientific). Alternatively, for Abca1 analyses, hemicortices of WT and *Mecp2* KO mice were homogenized in 1 mL of homogenization buffer (0.32 M sucrose, 1 mM Hepes, 0.1 mM EDTA) with protease and phosphatase inhibitor cocktail, by 30 strokes of glass-glass Potter homogenizer. The homogenates were centrifuged at 1,000g for 10 minutes at 4°C. Supernatants were collected and centrifuged at 9,000g for 15 minutes at 4°C and the pellets, corresponding to membrane-enriched fractions, were resuspended in a hypotonic buffer (20 mM Hepes, 0.1 mM DTT, 0.1 mM EDTA) with proteases and phosphatases inhibitor cocktails.

Samples containing 15-20 ug of proteins were prepared in sample buffer. For the analysis in the ACM, medium was mixed 1:1 with sample buffer. Samples were separated by SDS-PAGE on TGX-stain free precast gels (4-15% acrylamide gradient; #5678084, Bio-Rad) and proteins were blotted onto a nitrocellulose membrane using a semidry transfer apparatus (Trans-blot^®^ SD; Bio-Rad). Membranes were incubated 5 minutes in EveryBlot Blocking Buffer (#12010020, Bio-Rad) and then incubated O/N at 4°C with the primary antibody anti-Nsdhl (1:1000 in EveryBlot; #390871, Santa Cruz Biotechnology), anti-Abca1(1:1000 in EveryBlot; #PA1-16789, Invitrogen), anti-Lcat (1:1000 in EveryBlot; #bs1972R, Avantor) and anti-ApoE (1:800 in EveryBlot; #ab947, Merck). After three washes in Tris-buffered saline with Tween-20 (TBST), membranes were incubated with HRP-conjugated secondary antibody for 1 hours at RT (1:5.000; Jackson ImmunoResearch). The immunocomplexes were visualized by using the ECL substrate (Cyanagen and Bio-Rad) and Essential V6 imaging platform, UVITEC system (Cleaver Scientific Ltd). Band density measurements were performed using UVITEC software. Results were normalized to total protein content visualized by a TGX stain-free method.

### Non-denaturing gel electrophoresis

Concentrated ACM was mixed 1:1 with native sample buffer (#1610738, Bio-Rad) and run on a 4– 20% TGX-stain free precast gel in a Tris/Glycine Running buffer (#161-0734, Bio-Rad) at 100V at 4 °C. Gels were transferred to nitrocellulose membrane and probed with the anti-ApoE antibody (1:800 in EveryBlot). ApoE immunoreactivity was detected by chemiluminescent development with ECL reagents.

### Immunofluorescence staining and image analyses

Neurons and astrocytes were fixed in 4% paraformaldehyde (PFA), plus 10% of sucrose, for 8 minutes at RT. After three washes in PBS1X, cells were stored at 4°C with PBS1X containing 0.1% sodium azide (NaN3), until immunostaining. Cells were washed three times in PBS1X and then permeabilized in 0.2% Triton X-100 in PBS1X for 3 minutes on ice. After three washes in 0.2% of bovine serum albumin (BSA; #A3059; Merck) in PBS1X, cells were incubated in blocking solution (4% BSA in PBS1X) for 15 minutes at RT, followed by O/N incubation at 4°C with the primary antibodies diluted in PBS1X containing 0.2% BSA. For the analyses of synaptic puncta density, neurons were stained with antibodies against Synapsin1/2 (#106006, Synaptic System; 1:500), Shank2 (#162 211, Synaptic System; 1:300) and Map2 (#8707, Cell Signalling; 1:1000). An immunostaining with AnkyrinG (#386005, Synaptic System; 1:1000) was performed to detect axon initial segment (AIS) in cortical neurons. To assess Srebp2 expression and subcellular localization, astrocytes were stained with an antibody anti-Srebp2 (#LS-C179708, LS-Bio; 1:1000) and anti-Gfap (#MAB3402, Merck; 1:3000). To verify astrocyte purity following MACS isolation with magnetic beads, cells were stained with an antibody against an astrocyte marker (Gfap; #MAB3402, Merck; 1:3000), a microglial marker (Iba1; #019-19741, Wako; 1:1000), a neuronal marker (NeuN; #12943S, Cell Signalling; 1:1000) and an oligodendrocyte marker (Mbp; #AMAB91064, Merck; 1:1000). Alexa Fluor secondary antibodies were incubated for 1 hour at RT (1:500 in PBS containing 0.2% BSA). Nuclei were stained with DAPI (1:1.000 in PBS1X; #62248, ThermoFisher Scientific) for 5 minutes at RT, then cells were washed three times in PBS1X, and glass coverslips were mounted with Fluoromount-G™ Mounting Medium (#F4680, Merck) on microscope slides. For the detection of cholesterol in astrocytes, after immunostaining for Gfap, cells were incubated with filipin solution (#F4767, Merck powder in DMSO; 100 ug/ml in PBS1X) for 1 hour at RT in the dark. After 3 washes in PBS1X, DAPI staining was done, and glass coverslips were mounted.

A Leica Stellaris microscope equipped with an SP5 laser-scanning confocal system was used for the acquisition of synaptic puncta and Srebp2 immunofluorescence images. All images were acquired using identical acquisition parameters within each experiment.

*Synaptic puncta analysis*: synaptic puncta were quantified as previously described (Albizzati et al., 2024). Z-stack images (1024 × 1024 pixels, 8-bit grayscale) were acquired using a 63X oil-immersion objective at 1.4X digital zoom with a z-step of 0.3 μm. Synaptic puncta were quantified in Fiji (ImageJ) by analyzing three dendritic branches from at least 10 neurons per biological sample.

*Srebp2 immunofluorescence analysis:* cortical astrocytes were imaged using a 40X objective in a single focus plane. For each biological replicate, 5 randomly selected and non-overlapping fields of view were acquired. Srebp2 expression in cortical astrocytes was quantified by measuring the nuclear integrated density of Srebp2 immunofluorescence using Fiji software. Nuclear masks were generated from DAPI staining and superimposed onto the Srebp2 channel to extract the nuclear signal. Total Srebp2 expression was determined by measuring the overall integrated fluorescence intensity for each field and normalizing it to the number of cells.

A Nikon Eclipse Ti2-E widefield fluorescence microscope was used for the acquisition of AnkyrinG-immunostained neurons, filipin-stained astrocytes and acutely isolated astrocytes.

*Axon initial segment (AIS) analysis:* AIS length was quantified from AnkyrinG-stained neurons. An automatic threshold was applied to all images, and background fluorescence was calculated as the mean intensity of at least three regions of interest (ROIs) placed over cell nuclei. AIS length was determined by drawing a line along the axon and generating a fluorescence intensity profile. The AIS was defined as the portion of the profile in which the signal exceeded 33% of the normalized maximum fluorescence.

*Filipin analysis*: filipin fluorescence was imaged using UV excitation immediately after staining to minimize photobleaching. For each biological replicate, 5-10 randomly selected non-overlapping fields of view were acquired. Quantification was performed at the single-cell level by defining astrocyte boundaries using Gfap immunostaining. All images were acquired using identical microscope settings. Filipin fluorescence was measured as the integrated density within each Gfap-positive astrocyte and normalized to the corresponding astrocyte area.

*Assessment of astrocyte purity:* widefield fluorescence imaging was also used to evaluate the purity of acutely isolated astrocytes. Cell preparations were qualitatively assessed by immunofluorescence (IF) and showed no detectable neuronal or microglial contamination and only minimal contamination by oligodendrocytes (**Supplementary Figure 1**).

All image acquisition and quantitative image analyses were performed by investigators blinded to genotype.

### Cholesterol quantitative analysis

Cholesterol quantification was performed in WT and *Mecp2* KO cortical astrocytes, ACM and mouse cortical tissue using complementary analytical approaches.

*Gas chromatography-mass spectrometry (GC-MS) analysis of cholesterol and desmosterol:* intracellular cholesterol and desmosterol levels in cultured astrocytes, as well as cholesterol levels in ACM, were quantified by GC–MS. Briefly, samples were subjected to methyl *tert*-butyl ether extraction using 5-α-cholestane as an internal standard. The organic phase was collected after three sequential extraction steps, evaporated under nitrogen stream, dried under vacuum, and derivatized with *N,O*-bis(trimethylsilyl)trifluoroacetamide containing 1% trimethylchlorosilane for 2 hours at 60°C. GC-MS analyses were performed on a Trace GC-Ultra 60 gas chromatograph coupled to a Trace DSQ mass spectrometer (Thermo Scientific) equipped with a TraceGOLD TG5-MS capillary column. Cholesterol in ACM was quantified in selected ion monitoring (SIM) mode, whereas astrocyte extracts were analyzed in full-scan total ion current (TIC) mode, enabling cholesterol quantification together with the simultaneous detection of its biosynthetic intermediates. Under these analytical conditions, only cholesterol and desmosterol were detected and identified by comparison with authentic standards and published mass spectral data (Valverde-Som et al., 2018). All samples were analyzed in triplicate, and sterol levels were normalized to the total protein content of the corresponding samples.

*Amplex Red Cholesterol Assay:* AmplexRed Cholesterol Assay kit (#A12216, ThermoFisher Scientific) was used to measure cholesterol content in the ACM collected both from astrocytes derived from pups and adult (P40) mice, according to manufacturer instructions. Briefly, an enzymatic reaction was set up with 300 µM AmplexRed, 2 U/mL horseradish-peroxidase, 2 U/mL cholesterol oxidase, 0.2 U/mL cholesterol esterase in 1X reaction buffer. Samples were mixed 1:1 with reaction mix for a final volume of 100 µL and incubated at 37°C for 30 minutes in the dark. Fluorescence was measured with Spectramax i3x (Molecular Devices) plate-reader spectrophotometer at emission range of 550 ± 9 nm and excitation range of 590 ± 15 nm. Concentration of cholesterol was measured against a standard curve of cholesterol diluted serially in PBS1X reaction buffer.

*High Performance Thin Layer Chromatography (HPTLC) analysis:* cerebral cortexes were collected from WT and *Mecp2* KO mice and subjected to lyophilization. The extraction of total lipids from lyophilized tissue was carried out with the solvent system chloroform/methanol/water in proportion of 20:10:1 by vol (1.5 mL/50 mg dry tissue) mixing samples at 23°C in a thermomixer (Eppendorf) at 1,400 rpm for 15 minutes. Total lipid extract was separated from the pellet by centrifugation at 13,000g for 15 minutes, followed by a second and third extractions with chloroform/methanol, 2:1 by vol. Total lipid extracts were subjected to a two-phase partitioning by adding 20% water, resulting in the separation of an aqueous phase and an organic phase, used for cholesterol analysis. For each sample, equal amounts of organic phase corresponding to equal amounts of proteins were loaded on HPTLC (Silica gel 60 matrix on aluminum sheets with fluorescent indicator 254 nm) and cholesterol was separated by monodimensional HPTLC using hexane/ethylacetate (3:2, v/v) solvent system. After lipid separation, HPTLC was revealed using the anisaldehyde reagent and cholesterol was identified and quantified by co-migration with dosed cholesterol standards. HPTLC was scanned and the band intensity quantified using the ImageJ software (2.14.0; Java 1.8.0_322, NIH, Bethesda, MD, USA; http://rsbweb.nih.gov/ij/ accessed on 1 July 2023); each band was normalized to the total protein content.

### Cholesterol treatment on neurons

Cholesterol complexed with methyl-β-cyclodextrin (MβCD) was used for all supplementation experiments (#C4951, Merck). A starting concentration of a cholesterol solution (20 μg/ml in culture medium) was added on DIV13 primary neurons to obtain a final concentration of 0.1 μg/ml. Cholesterol treatment last 24 hours. Untreated neurons were supplemented with an equal volume of neuron culture medium.

### Statistical analysis

All statistical analyses were performed using Prism 9 (GraphPad Software). Data are expressed as mean ± SEM, except for violin plots in which the bar indicates the median with interquartile range. For each dataset, D’Agostino-Pearson and Shapiro-Wilk tests were used to evaluate normality distribution, and depending on the result, either the unpaired Student’s t-test or the Mann-Whitney test were chosen for two groups’ comparison analysis. Multiple groups’ comparisons were performed through 2-way Anova, followed by Tukey’s post hoc test. Before each statistical analysis, outliers were assessed using ROUT test (Q = 1%) or Grubbs’ test (α = 0.05%) and then excluded. A p-value <0.05 was considered significant. *p<0.05, **p<0.01, ***p<0.001.

## Results

### Cholesterol metabolic pathway is impaired in cortical *Mecp2* KO astrocytes

Astrocytes are the main source of cholesterol in the brain, and cholesterol secreted by astrocytes has a role in synaptic formation and maintenance (Goritz et al., 2005; J.-P. Liu et al., 2010; Pfrieger, 2003). Considering the defective capacity of RTT astrocytes to support synaptogenesis and synaptic functionality (Albizzati et al., 2024; Sun et al., 2023), we characterized cholesterol metabolism in *Mecp2* deficient astrocytes. We first investigated the subcellular localization of sterol regulatory element-binding protein 2 (Srebp2), one of the master transcriptional regulators of cholesterol homeostasis. Immunofluorescence (IF) analysis revealed a significant reduction in nuclear Srebp2 staining in KO astrocytes compared with WT cells (**Figure 1A,C**), indicating impaired translocation to nucleus, the compartment where Srebp2 is transcriptionally active. In contrast, total Srebp2 protein abundance and mRNA expression were unchanged (**Figure 1B, Supplementary Figure 2**), suggesting that *Mecp2* deficiency affects Srebp2 activation rather than its expression. Given the reduced nuclear localization of Srebp2, we next investigated the expression of genes involved in cholesterol metabolism. In primary cortical astrocytes isolated from P1–P3 mice, qRT-PCR analysis revealed a broad downregulation of genes involved in both cholesterol biosynthesis and transport in KO cells compared with WT. Specifically, transcripts encoding key enzymes of the cholesterol biosynthetic pathway, including Nsdhl, Hmgcr, Mvk and Sqle, were significantly reduced. Similarly, the expression of Abca1, Abcg1 and Lcat, which participate in extracellular cholesterol transport, was markedly decreased. In contrast, ApoE mRNA levels were not altered (**Figure 1D**). To determine whether these transcriptional changes were also detectable in acutely sorted astrocytes, we isolated cortical astrocytes from P7 WT and KO mice by magnetic beads, providing validation in a serum-free astrocyte preparation. Consistent with the findings obtained in pups-derived astrocyte cultures, qRT-PCR analysis confirmed the downregulation of most cholesterol-related genes in KO astrocytes compared with WT cells (**Figure 1E**).

**Figure 1.**
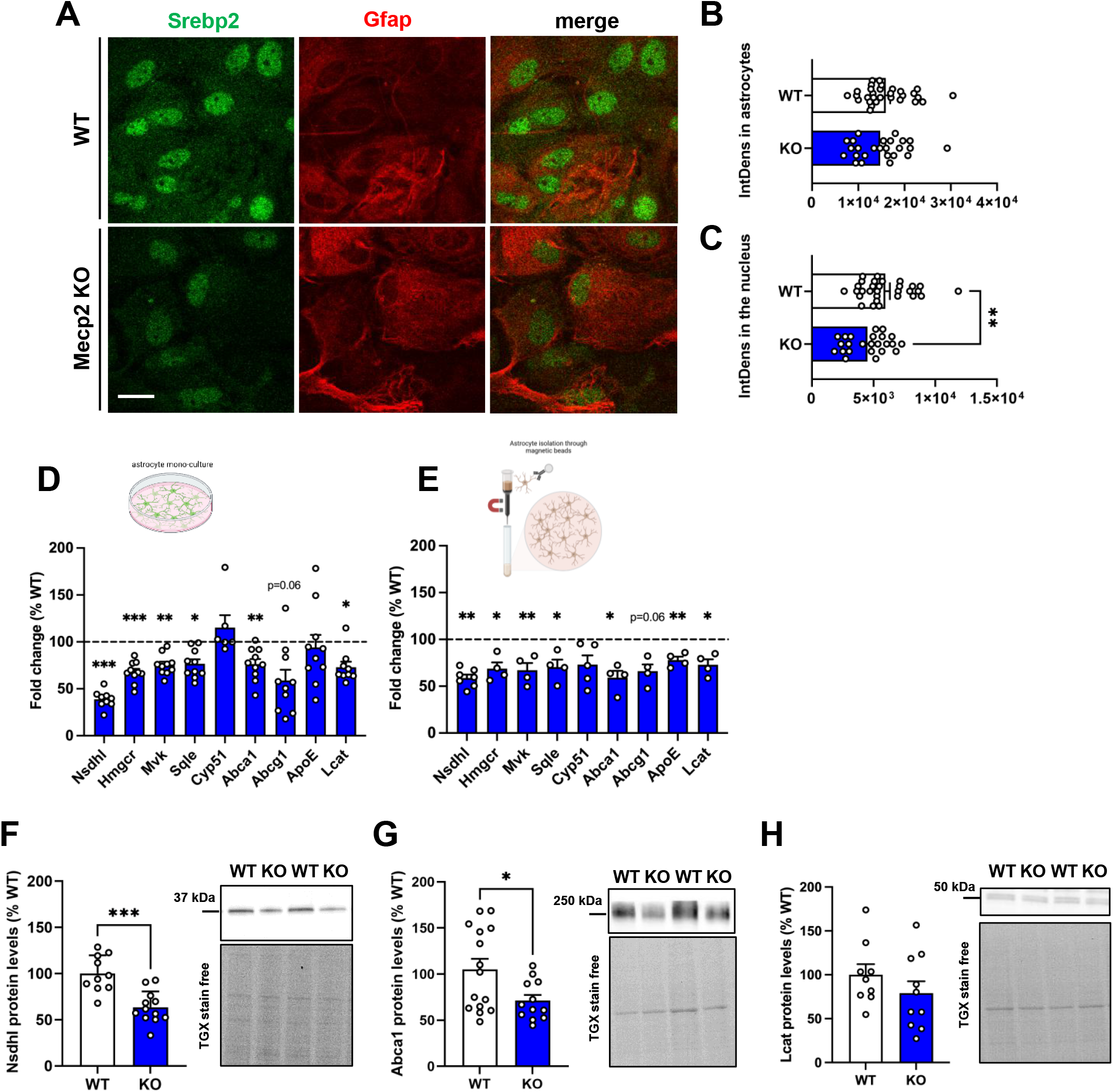
Molecular deregulation of the expression of cholesterol-related genes in *Mecp2* KO astrocytes. (**A**) Representative immunofluorescence images of Srebp2 (green) and Gfap (red) in WT and *Mecp2* KO cortical astrocytes, along with the merged signals. Scale bar=20µm. (**B,C**). Histograms report the integrated density measured in the whole astrocytes (B) and selectively in the nucleus (C). Data are shown as mean±SEM. n=25/30 fields deriving from n=6/7 biological replicates. **p<0.01 by Student’s t-test. (**D,E**) Histograms show the mRNA levels of genes involved in the synthesis and transport of cholesterol in cortical astrocytes isolated from P1-P3 pups (D) and in astrocytes isolated through MACS technology from the cortex of P7 mice (E). In both graphs, data are expressed as percentages with respect to WT astrocytes, set at 100% and represented by the dotted line. Data are shown as mean±SEM. n=6/10. Statistical analysis was performed by Student’s t-test or Mann-Whitney test according to data distribution; *p<0.05, **p<0.01, ***p<0.001. (**F-H**) Histograms depict the protein levels of Nsdhl (F), Abca1 (G) and Lcat (H) in *Mecp2* KO astrocytes, compared to WT astrocytes. Data, expressed as percentages with respect to WT astrocytes, are shown as mean±SEM. n= 6/exp.group. Statistical analysis was performed by Student’s t-test or Mann-Whitney test according to data distribution: *p<0.05, ***p<0.001. Representative bands of Nsdhl, Abca1 and Lcat and the total protein content, visualized by TGX stain-free technology, are shown.

We next assessed the protein levels of selected cholesterol-related proteins by western blot (WB). Consistent with the mRNA data, Nsdhl protein expression was markedly reduced in KO astrocytes compared with WT cells (**Figure 1F**). Similarly, Abca1 (ATP-binding cassette transporter A1), a key mediator of ApoE lipidation, was significantly decreased in *Mecp2* KO astrocytes (**Figure 1G**), whereas Lcat (lecithin–cholesterol acyltransferase), which contributes to the maturation of ApoE-containing lipoprotein particles, showed only a trend toward reduced expression (**Figure 1H**).

### ApoE lipidation is defective in cortical *Mecp2* KO astrocytes

To determine whether the observed changes in gene expression translated into altered cholesterol synthesis and secretion, we quantified cholesterol levels both in astrocytes and astrocyte-conditioned medium (ACM) derived from WT and *Mecp2* KO cultures (**Figure 2**). Gas chromatography-mass spectrometry (GC-MS) analysis revealed a trend toward increased cholesterol levels in KO astrocytes, together with a significant accumulation of its precursor desmosterol (**Figure 2A**). Consistent with the GC-MS data, Filipin staining revealed an increased accumulation of intracellular free (unesterified) cholesterol in KO astrocytes. Quantification of the integrated fluorescence density per cell confirmed a significant accumulation of free intracellular cholesterol compared with WT astrocytes (**Figure 2B,C**). Notably, the accumulation of intracellular cholesterol provides a potential explanation for the reduced nuclear localization of Srebp2 observed in KO astrocytes, given the negative feedback exerted by intracellular cholesterol on Srebp2 activation (Duan et al., 2022; N.-Q. Wang et al., 2025).

**Figure 2.**
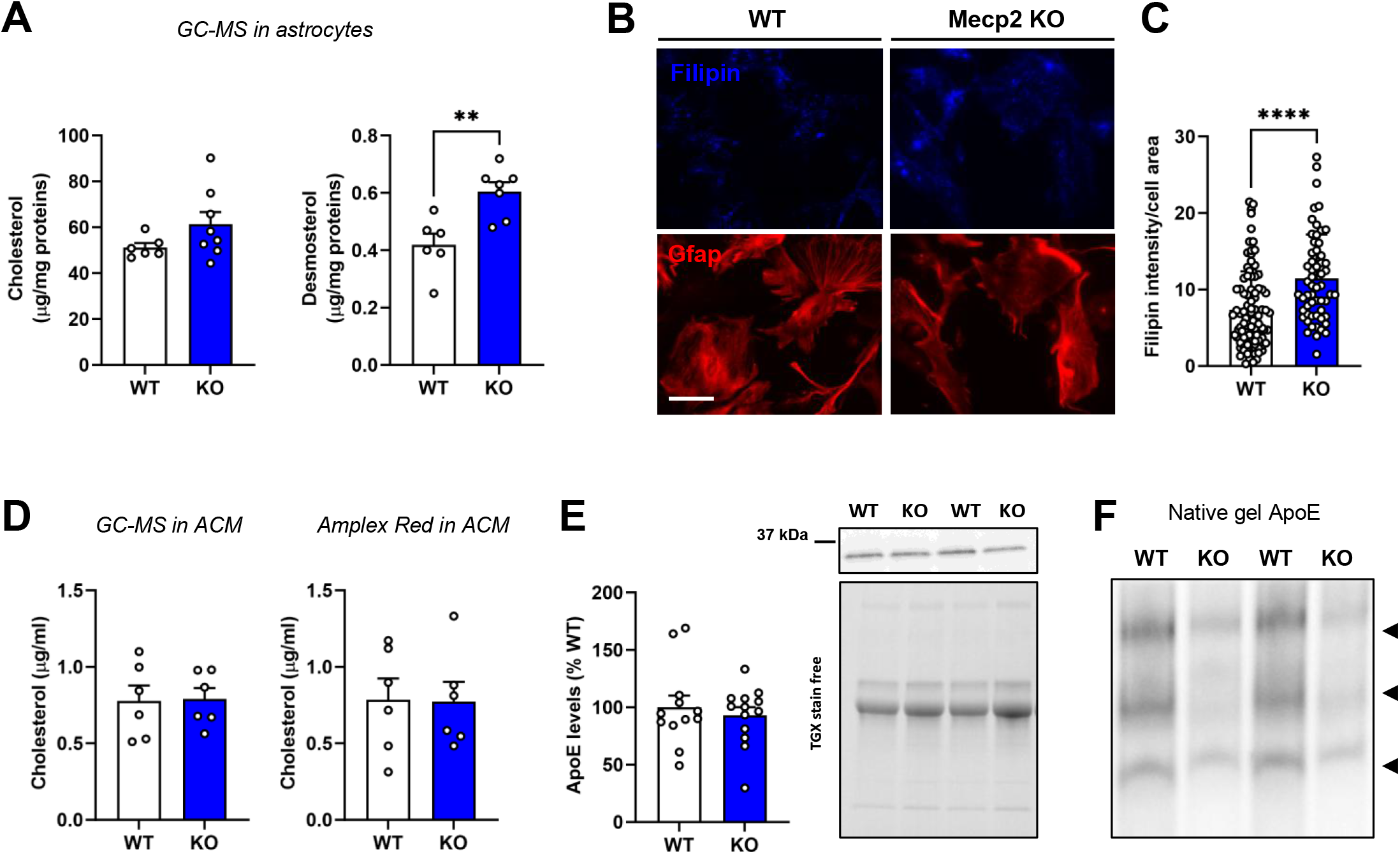
Mecp2 deficiency does not affect astrocytic cholesterol secretion but impairs ApoE lipidation. **(A)** Histograms show the content of cholesterol and its precursor desmosterol in WT and *Mecp2* KO cortical astrocytes analysed through GC-MS and normalized to protein concentration. Data are expressed as mean±SEM, n=6-8/exp.group. **p<0.01, by Mann-Whitney test. **(B)** Representative images of Filipin (blue) and Gfap (red) in WT and *Mecp2* KO cortical astrocytes. Scale bar=20µm. (**C**) The graph shows the integrated density of Filipin signal within the area of the astrocytes. n=80/100. Data are shown as mean±SEM. ****p<0.0001 by Student’s t-test. (**D**) Graphs depict cholesterol levels in the medium of WT and *Mecp2* KO cortical astrocytes obtained from P1-P3 mice and measured through GC-MS and Amplex Red assay. In both cases, cholesterol concentration in the medium was normalized to the protein concentration of cells; n=6/exp.group, deriving from at least 2 independent experiments. Statistical analysis was performed with the Mann-Whitney test. **(E)** The graph shows ApoE protein levels in the medium of *Mecp2* KO astrocytes, expressed as percentage of WT astrocytes; n=12/exp.group. Statistical analysis was performed by Student’s t-test. Representative bands of ApoE and the total protein content, visualized by TGX stain-free technology, are shown. **(F)** Native gel electrophoresis probed for ApoE, showing differences in the lipidation of ApoE in the medium of *Mecp2* KO astrocytes compared to WT.

Despite the increase in intracellular cholesterol, it remained unclear whether cholesterol released by KO astrocytes, the fraction directly involved in astrocyte-neuron communication, was also affected. We therefore quantified cholesterol levels in ACM. GC-MS reported no alteration in cholesterol concentration in the KO-ACM, with respect to WT secretome, and similar values resulted from the enzymatic fluorometric Amplex Red Cholesterol Assay (**Figure 2D**). The amount of cholesterol secreted by KO astrocytes did not change even when the analysis was extended to the ACM of acutely isolated astrocytes, thus excluding the possibility that a lack of difference was attributed to the protocol used (**Supplementary Figure 3**). Because the functional efficacy of cholesterol relies on its correct trafficking to neurons via apolipoproteins, we considered it important to analyse both ApoE levels as well as its capacity to complex cholesterol. Through WB, we proved that ApoE expression remained unaffected in the ACM collected from KO astrocytes (**Figure 2E**). However, analysis of its lipidic state, qualitatively assessed by an electrophoresis in non-denaturating native gel, indicated a strong impairment, as reported by an evident reduction of the levels of lipidated ApoE complexes in KO-ACM with respect to WT-ACM (**Figure 2F**). Of relevance, the reduction in ApoE lipidation in KO samples might be ascribed to the reduction of Abca1 transporter, considering its key role in free cholesterol transfer to ApoE (C. Liu et al., 2025) (Figure 1).

### Molecular evidence of dysfunctional cholesterol metabolism in the *Mecp2* deficient brain

To corroborate *in vitro* evidence, we first quantified cholesterol levels in the cerebral cortex of symptomatic (P40) *Mecp2* KO mice. Consistent with the results obtained in cultured astrocytes and previous evidence (Buchovecky et al., 2013a), High Performance Thin Layer Chromatography (HPTLC) analysis revealed a significant increase in cortical cholesterol levels in KO mice compared with WT controls (**Figure 3A, Supplementary Figure 4).** We next investigated whether this phenotype was accompanied by the transcriptional alterations observed in cultured astrocytes. Gene set enrichment analysis (GSEA) of our previously published cortical transcriptomic dataset (Pozzer et al., 2025) revealed a significant impairment of cholesterol homeostasis pathway in *Mecp2* KO cortex (**Figure 3B**). By validating the reduced expression of selected cholesterol-related genes, qRT-PCR revealed a significant reduction of Nsdhl, Hmgcr, Sqle and Cyp51, and a trend toward a reduction for Mvk in the whole cerebral cortex of symptomatic (P40) *Mecp2* KO mice (**Figure 3C)**. In addition, our previous study reported a significant down-regulation of Lcat in the *Mecp2* KO cortex (Albizzati et al., 2022). To investigate whether these defects were attributable to astrocytes, we performed gene expression analyses on cortical astrocytes acutely sorted from adult (P40) mice with anti-ACSA-2-conjugated magnetic beads. We observed a consistent downregulation of Nsdhl, Hmgcr and Cyp51, as well as a trend toward reduced expression of Abcg1 and Lcat in KO cells, further supporting the existence of a general impairment of cholesterol homeostasis (**Supplementary Figure 5**).

**Figure 3.**
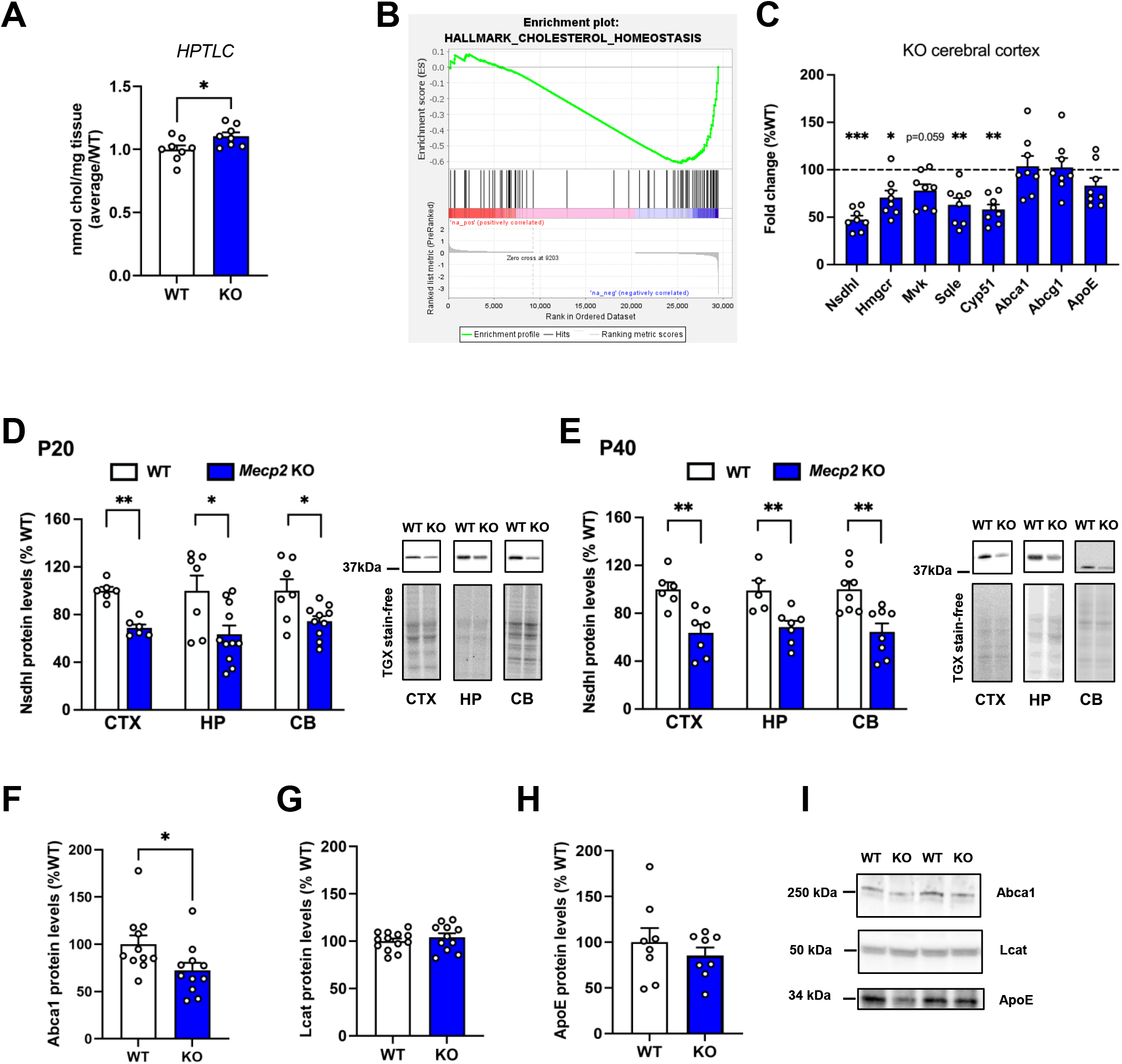
Molecular evidence of cholesterol metabolism deregulation in the cerebral cortex of Mecp2 deficient animals. **(A)** Graph shows the content of cholesterol in the cortex of *Mecp2* KO (P40) mice and age-matched WT littermates analysed through HPTLC. Data are normalized on protein content. n=8/exp.groups *p<0.05 by Student’s t test. In the graph, data are expressed relative to WT. **(B)** Gene set enrichment analysis (GSEA) of cholesterol homeostasis using RNA-sequencing data from the cortex of WT and *Mecp2* KO P60 mice. NES = -1.57 and FDR q=0.044. RNA sequencing data deriving from (Pozzer et al., 2025). **(C)** Graph shows the mRNA levels of genes involved in the synthesis and transport of cholesterol in the cortex of symptomatic *Mecp2* KO (P40). Data are expressed as percentages with respect to male WT littermates, set at 100% and represented by the dotted line. Data are shown as mean±SEM. n=8/exp. groups.; *p<0.05, **p<0.01, ***p<0.001 by Student’s t test. (**D-E**) Histograms show the protein expression levels of Nsdhl measured in the cortex (CTX), hippocampus (HP) and cerebellum (CB) of pre-symptomatic (at P20) (D) and symptomatic (P40) (E) *Mecp2* KO animals. Data are expressed as percentages of corresponding WT animals and shown as mean±SEM. n=5-10/exp.groups. *p<0.05, **p<0.01 by Student’s t test. On the right of each graph, representative images of Nsdhl protein and total protein content visualized by TGX stain-free technology are shown. **(F-H)** Histograms report the protein expression level of Abca1(F), Lcat (G) and ApoE (H) in the cortex of WT and KO mice at P60. Data are expressed as percentages of corresponding WT animals and shown as mean±SEM. n=8-13/exp.group. *p<0.05 by Student’s t-test or Mann-Whitney test according to data distribution, (**I**) Representative images of Abca1, Lcat and ApoE protein bands are reported.

By WB, we reported a marked reduction in the expression of Nsdhl, not only in the cerebral cortex but also in the hippocampus and cerebellum. Notably, these decrements were already evident in KO animals at the pre-symptomatic stage (P20) and maintained at the symptomatic one (P40) (**Figure 3D,E**). Importantly, Nsdhl protein levels were similarly reduced by approximately 50% in the brain of the KI Mecp2^Y120D/y^ mouse model, which carries a phospho-mimetic missense mutation and phenotypically recapitulates the null mouse **(Supplementary Figure 6A)** (D’Annessa et al., 2018; Gandaglia et al., 2019). In contrast, Nsdhl downregulation in symptomatic *Mecp2* HET female mice (P200) was restricted to the cortex (**Supplementary Figure 6B**). To investigate whether ApoE-mediated cholesterol trafficking was also impaired in the mouse brain, we analysed the protein expression of ApoE, Abca1 and Lcat. Obtained results indicated that, whereas ApoE and Lcat protein levels were not affected, a significant reduction of Abca1 was evident in the cortex of KO animals (**Figure 3F-I**).

### Cholesterol supplementation prevents synaptic defects in WT neurons induced by *Mecp2* KO Astrocyte Conditioned Medium and in *Mecp2* heterozygous neurons

To assess whether reduced cholesterol trafficking to neurons plays a role in the occurrence of the synaptic defects observed in WT neurons treated with KO-ACM, we found it interesting to investigate whether supplementation of KO-ACM with exogenous cholesterol could rescue synaptic alterations. For these experiments, we used a water-soluble cholesterol formulation complexed to methyl-β-cyclodextrin, that has been already used in similar experimental settings because of its solubility and ability to be internalized into cells (Korinek et al., 2020; Marquer et al., 2014; Moutinho et al., 2016). We initially selected a non-toxic dose, by testing in WT cortical neurons the safety of increasing concentrations of the water-soluble formulation of cholesterol, added at DIV13 for 24 hours. The dosage of 0.1 μg/ml was thus supplied to KO-ACM, which was used to treat WT neurons (from DIV13 to DIV14). As control, WT neurons were treated with WT-ACM in the presence of or absence of cholesterol. IF analysis for pre-synaptic and post-synaptic markers confirmed the already reported defects in neurons treated with KO-ACM (Albizzati et al., 2024) and demonstrated that the addition of cholesterol rescues synaptic alterations. In particular, the reduced density of both pre- and post-synaptic compartments, as well as that of colocalized puncta, was fully reversed by cholesterol treatment (**Figure 4A-D**).

**Figure 4.**
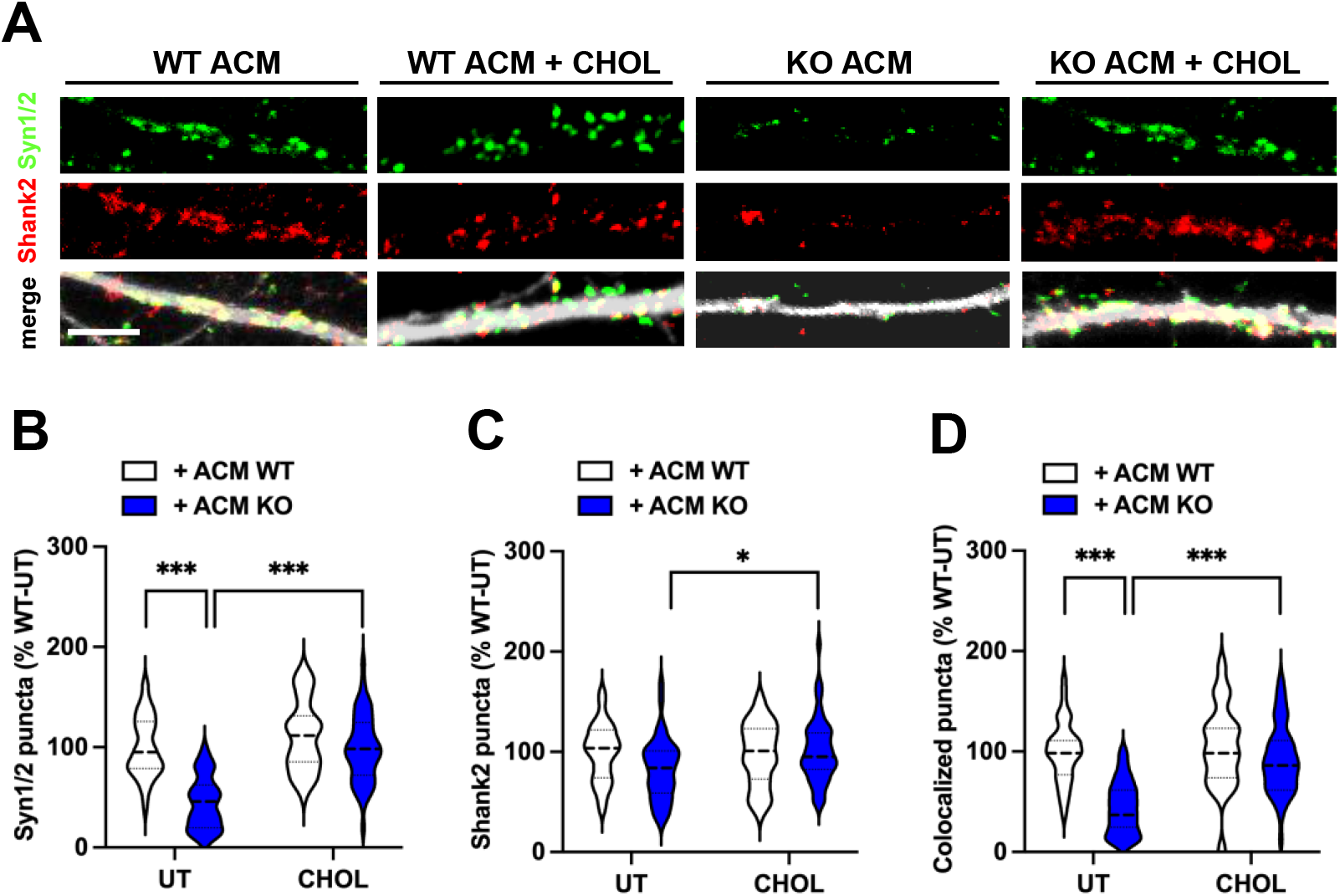
Cholesterol treatment rescues synaptic alterations induced by *Mecp2* KO-ACM. (**A**) Representative images of primary branches from WT neurons treated for 24 hours with WT-ACM or KO-ACM, supplemented or not with cholesterol (0.1 μg/ml), and immunostained for Synapsin1/2 (Syn1/2; green), Shank2 (red) and Map2 (white). Scale bar=5 µm. (**B-D**) Violin plots indicate the median (dashed line) and 25^th^ and 75^th^ percentiles (dotted lines) of Synapsin1/2 (B), Shank2 (C) and colocalized puncta number (D). Values for puncta number are expressed as percentages with respect to neurons treated with WT-ACM. n=40/50. Statistical analysis was performed by 2-way Anova, followed by Sidak post-hoc test; *p<0.05, ***p<0.001.

To verify whether a similar synaptogenic effect of cholesterol was also evident in *Mecp2* HET neurons, the same paradigm of cholesterol treatment (0.1 μg/ml) was applied on neurons for 24 hours (from DIV13 to DIV14). By measuring the density of synaptic puncta, we proved that cholesterol treatment recovers pre- and post-synaptic defects, typically shown by HET neurons (Frasca et al., 2024) **(Figure 5A-D)**.

**Figure 5.**
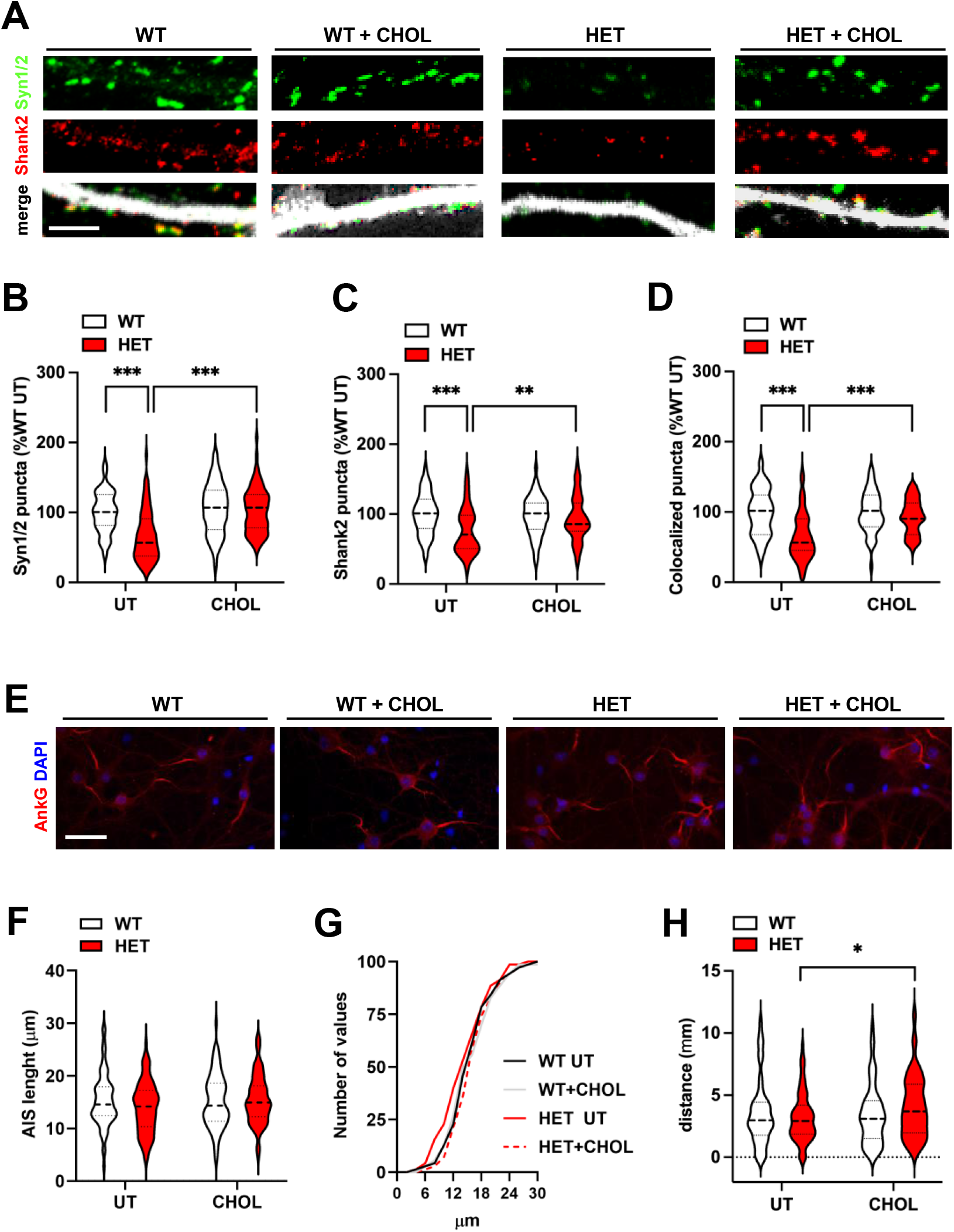
Cholesterol treatment rescues synaptic alterations in *Mecp2* HET neurons. (**A**) Representative images of primary branches from WT and *Mecp2* HET neurons, treated for 24 hours with cholesterol or with vehicle (0.1 μg/ml), and immunostained for Synapsin1/2 (Syn1/2; green), Shank2 (red) and Map2 (white). Scale bar=5 µm. (**B-D**) Violin plots indicate the median (dashed line) and 25^th^ and 75^th^ percentiles (dotted lines) of Synapsin1/2 (B), Shank2 (C) and colocalized puncta number (D). Values for puncta number are expressed as percentages with respect to untreated WT neurons. n=57/69. Statistical analysis was performed by 2-way Anova, followed by Sidak post-hoc test; **p<0.01, ***p<0.001. (**E**) Representative images of axon initial segment (AIS) stained with anti-AnkyrinG (AnkG; red). Scale bar=20 µm. (**F,G**) Violin plots indicate the median (dashed line) and 25^th^ and 75^th^ percentiles (dotted lines) of AIS length in WT and *Mecp2* HET neurons (at DIV14) after cholesterol supplementation or treated with vehicle (F). The frequency distribution of values of AIS length is shown in (G). n=70/exp.groups. (**H**) Violin plots indicate the median (dashed line) and 25^th^ and 75^th^ percentiles (dotted lines) of the distance of AIS from the soma in WT and *Mecp2* HET neurons (at DIV14) after cholesterol supplementation or treated with vehicle. n=70/exp.groups. *p<0.05, by 2-way Anova, followed by Sidak post-hoc test;

To investigate the effects of cholesterol treatment on neuronal function, we measured the length of the axon initial segment (AIS), a key structure responsible for action potential generation. AIS length was assessed by immunostaining for the scaffold protein Ankyrin-G (Hedstrom et al., 2008; Ogawa & Rasband, 2008) (**Figure 5E**). Image analysis revealed no significant differences in mean AIS length between HET cortical neurons and WT cells, neither in basal conditions nor following cholesterol treatment (**Figure 5F**). However, analysis of the frequency distribution showed an increase in the percentage of HET neurons exhibiting shorter AIS compared to WT cells, a phenotype that was not observed after cholesterol supplementation. Specifically, we estimated that approximately 40% of HET neurons displayed an AIS shorter than 12 µm, compared to about 20% of the treated experimental groups (**Figure 5G**). By analyzing AIS positioning relative to the soma, no significant differences were detected between untreated HET and WT neurons, whereas cholesterol treatment induced a significant distal shift of the AIS in HET neurons, suggesting an activity-dependent remodeling of the AIS associated with the restoration of synaptic function (**Figure 5H**).

## Discussion

Synaptic defects are a hallmark of RTT (Bedogni et al., 2014; Boggio et al., 2010; Calfa et al., 2015; Dani et al., 2005; Fukuda et al., 2005) and *MECP2* mutant astrocytes contribute to neuronal alterations through non-cell-autonomous mechanisms (Albizzati et al., 2024; Ballas et al., 2009; Sun et al., 2023; Williams et al., 2014). Indeed, through the release of synaptotoxic molecules and/or the reduced secretion of synaptogenic factors, *Mecp2* KO astrocytes influence both synaptogenesis and synapse maintenance (Albizzati et al., 2024; Sun et al., 2023). After demonstrating the excessive secretion of Interleukin-6 by KO astrocytes and its detrimental role for synapses (Albizzati et al., 2024), here we have investigated the role of cholesterol in the astrocyte-neuron crosstalk in RTT. This choice was driven by the well-established role of cholesterol in synaptic development and maintenance (Goritz et al., 2005; J.-P. Liu et al., 2010; Pfrieger, 2003), as well as by accumulating evidence of altered brain cholesterol metabolism in RTT (Ben-Shachar et al., 2009; Buchovecky et al., 2013b; Lopez et al., 2017; Lütjohann et al., 2018; Urdinguio et al., 2008; Villani et al., 2016; Zandl-Lang et al., 2022). Several data reported dysfunctions in cholesterol metabolism in both RTT murine models and human samples. In particular, *Mecp2* KO mice exhibit a reduced rate of brain cholesterol biosynthesis, together with the downregulation of key metabolic enzymes and decreased levels of cholesterol precursors, despite the accumulation of cholesterol in the brain (Ben-Shachar et al., 2009; Buchovecky et al., 2013b; Lütjohann et al., 2018; Urdinguio et al., 2008, 2008; Villani et al., 2016). Moreover, reduced cholesterol levels have been recently reported in the cerebrospinal fluid of RTT patients (Zandl-Lang et al., 2022). Despite these findings, the relevance of cholesterol metabolism in astrocyte-neuron communication has not been previously addressed, particularly with respect to the metabolically active pool of cholesterol involved in intercellular trafficking, which is functionally distinct from cholesterol present in myelin, cellular membranes and in intracellular stores.

Our molecular analyses of *Mecp2* KO cortical astrocytes revealed a widespread downregulation of genes involved in cholesterol biosynthesis and transport, suggesting impairments in both cholesterol production and its extracellular delivery. Consistent with these transcriptional changes, we found reduced nuclear localization of Srebp2, one of the master transcriptional regulators of cholesterol metabolism, indicating impaired Srebp2 activation in KO astrocytes. The reduced expression of cholesterol biosynthetic genes appears paradoxical given the intracellular accumulation of cholesterol detected in KO astrocytes. However, intracellular cholesterol is known to inhibit Srebp2 activation through a negative feedback mechanism (Duan et al., 2022; Wang et al., 2025). Thus, cholesterol accumulation itself might account for the reduced nuclear translocation of Srebp2 and, consequently, for the widespread transcriptional inactivation of its target genes. Alternatively, gene deregulation might also arise from the loss of *Mecp2*-dependent transcriptional control. While this mechanism is unlikely to involve direct regulation of Srebp2, whose mRNA levels remained unchanged, other cholesterol-related genes might represent direct MeCP2 targets. Notably, genes such as Nsdhl were consistently downregulated across different experimental models, supporting the existence of a MeCP2-dependent transcriptional program controlling a subset of cholesterol-related genes. Interestingly, Nsdhl downregulation has also been reported in an independent study of Rett syndrome (Luoni et al., 2020), further supporting the reproducibility of this finding. This observation is particularly intriguing considering that pathogenic variants in *NSDHL*, which cause CHILD syndrome (Congenital Hemidysplasia with Ichthyosiform nevus and Limb Defects), lead to defective cholesterol biosynthesis and severe developmental abnormalities, including neurological manifestations (König et al., 2000). Whether *NSDHL* represents a direct MeCP2 transcriptional target and whether its reduced expression is sufficient to impair cholesterol homeostasis in astrocytes will require further investigation.

Importantly, cholesterol metabolism alterations were not restricted to cultured astrocytes. Similar changes in the expression of cholesterol-related genes were detected in acutely isolated cortical astrocytes and in the cerebral cortex from *Mecp2* KO mice, indicating that the dysregulation of cholesterol homeostasis is also present *in vivo*. Moreover, the increased cholesterol levels detected in the cerebral cortex of symptomatic *Mecp2* KO mice further support the presence of altered sterol homeostasis in the diseased brain. Although these findings obtained on tissues were limited to the quantification of cholesterol and the assessment of gene and protein expression, future studies will investigate the broader lipidomic profile of *Mecp2* deficient brains.

One of the most relevant findings of this study is the marked reduction in ApoE lipidation in *Mecp2* KO astrocytes, indicating that, despite preserved cholesterol secretion, the fraction of cholesterol that is released in a functional lipoprotein-associated form and available for delivery to neurons is significantly reduced. Notably, this defect occurred despite unchanged ApoE expression, suggesting that Mecp2 deficiency primarily impairs ApoE maturation rather than its synthesis. A plausible explanation for reduced ApoE lipidation is the decreased expression of Abca1, the major cholesterol transporter responsible for the initial lipidation of ApoE (Liu et al., 2025), which we observed both in cultured astrocytes and in the cerebral cortex of KO mice. Reduced Abca1 expression is expected to limit cholesterol and other lipids loading onto ApoE particles, thereby impairing the formation of mature lipoproteins. Although the present study does not establish a direct causal link between impaired ApoE lipidation and synaptic dysfunction, our findings strongly support the hypothesis that defective ApoE maturation limits cholesterol bioavailability to neurons and contributes to the observed synaptic phenotype

Collectively our findings suggest that the primary defect in RTT might not reside in the total amount of cholesterol present in the brain, but rather in its functional availability and trafficking to neurons. Indeed, total cholesterol levels do not necessarily reflect the fraction of cholesterol that is effectively available for astrocyte-to-neuron transport, a concept that should be considered when designing therapeutic strategies targeting cholesterol metabolism. Accordingly, pharmacological strategies aimed at lowering total cholesterol levels, including statins, might not adequately address the underlying defect and could even prove ineffective or detrimental, as supported by results obtained from both preclinical studies and a phase II clinical trial in RTT patients (Villani et al., 2016). Our findings support the idea that restoring cholesterol trafficking and its functional delivery to neurons, rather than simply reducing cholesterol levels, might represent a more appropriate therapeutic strategy.

Consistent with this hypothesis, bypassing the defective astrocyte-derived cholesterol transport through exogenous cholesterol supplementation fully rescued the synaptic defects induced in WT neurons by factors secreted from *Mecp2* KO astrocytes. Notably, a similar beneficial effect was observed in *Mecp2* HET neurons, a model that more closely recapitulates the mosaic condition of RTT patients. Moreover, the restoration of AIS length together with the distal repositioning of the AIS following cholesterol treatment suggests that cholesterol not only rescues synaptic structural defects but also restores activity-dependent mechanisms regulating intrinsic neuronal excitability. The beneficial effects of cholesterol supplementation are likely to rely on the multiple functions of cholesterol in synaptic physiology. Beyond serving as a structural component of cellular membranes, cholesterol is essential for synapse formation and maintenance. Indeed cholesterol depletion has been associated with dendritic spine loss and synaptic degeneration, whereas its supplementation promotes synapse formation and enhances synaptic activity (Goritz et al., 2005; Liu et al., 2010; Mauch et al., 2001). One mechanism that might contribute to these effects is the restoration of membrane lipid rafts, cholesterol-enriched microdomains that organize signaling molecules and synaptic proteins and are required for efficient synaptic transmission (Korinek et al., 2020; Linetti et al., 2010). To our knowledge, direct evidence for lipid raft alterations in RTT is currently lacking. Nevertheless, alterations in lipid raft composition and function have also been implicated in autism spectrum disorders (Wang, 2014) and several signaling pathways that are disrupted in RTT, including BDNF/TrkB signaling, glutamatergic neurotransmission and activity-dependent synaptic plasticity, are known to depend on the integrity of lipid rafts for their proper organization and function (Sebastião et al., 2013; Zonta & Minichiello, 2013). The therapeutic relevance of targeting cholesterol homeostasis is further supported by a recent gene therapy study, reporting that restoring Cyp46a1 expression in *Mecp2* KO mice significantly ameliorated disease manifestations (Audouard et al., 2024), providing independent evidence that improving cholesterol homeostasis leads to behavioral benefits.

Our findings are consistent with growing evidence implicating astrocyte-dependent cholesterol dysregulation in several neurological disorders, including Fragile X syndrome, Alzheimer’s disease and Huntington’s disease (Ren et al., 2023; Valenza et al., 2010). Our study extends these observations to RTT by identifying impaired cholesterol trafficking as a potential driver of synaptic dysfunction. Together with previous reports demonstrating the beneficial effects of cholesterol supplementation in other neurological disorders (Birolini et al., 2023; Valenza, 2025) our findings support strategies aimed at restoring cholesterol availability to neurons as a promising therapeutic avenue for RTT.

In conclusion, our study identifies impaired cholesterol trafficking from astrocytes to neurons as a novel mechanism contributing to synaptic dysfunction in RTT and highlights cholesterol metabolism as a potential therapeutic target for this disorder.

## Supporting information

Supplementary results

## Acknowledgements

This work was supported by the Italian parents’ association “ProRETT Ricerca” to A.F. F.M.P salary is supported by the PhD Course in Experimental Medicine of the University of Milan.

## Conflict of interest

The authors declare that they have not conflict of interest.

## Notes

### Competing Interest Statement

The authors have declared no competing interest.

