## Supplementary results for "Impaired astrocyte-to-neuron cholesterol trafficking drives synaptic dysfunction in Rett syndrome"

**Supplementary Figures**

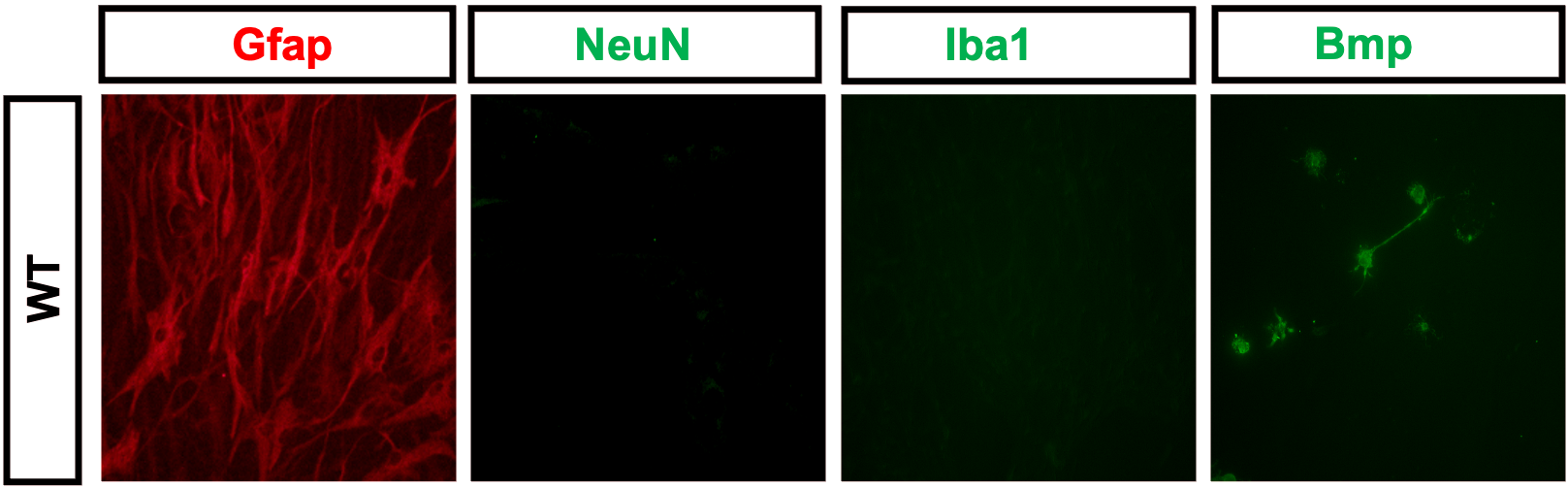

**Figure S1.** Representative images of MACS-isolated astrocytes from the cortex of P7 animals stained for the astrocytic marker Gfap, the neuronal marker NeuN, the microglial marker Iba1 and the oligodendrocytic marker Bmp, to assess astrocyte purity.

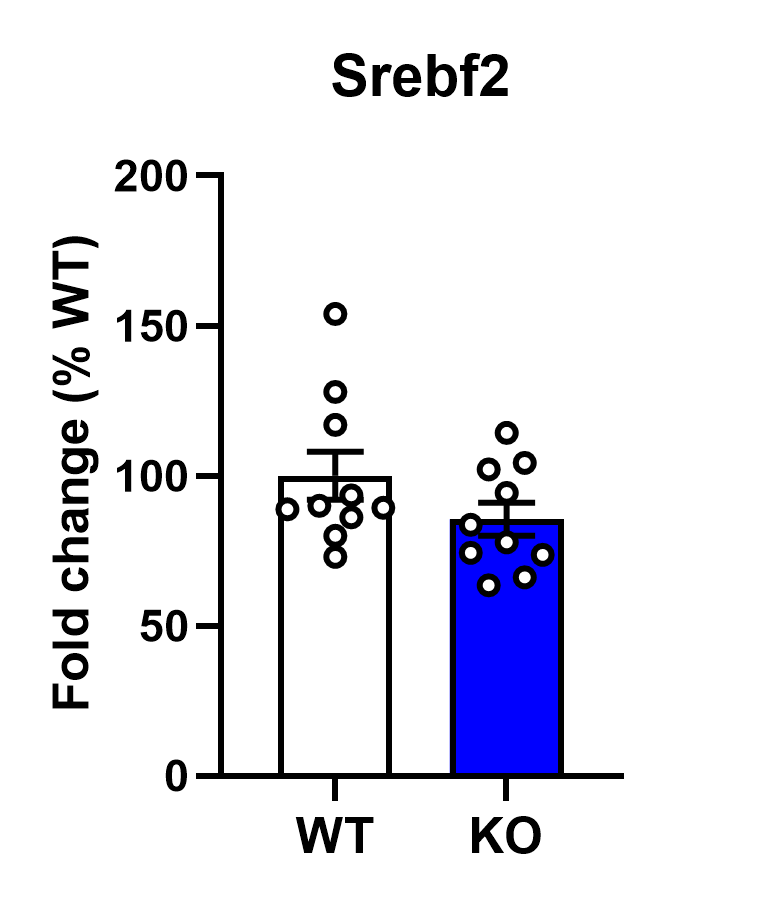

**Figure S2.** Histogram shows the the mRNA levels of Srebf2 in WT and *Mecp2* KO cortical astrocytes. Statistical analysis was performed by Student’s t test. n=10/exp.groups.

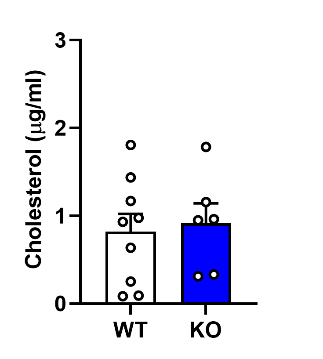

**Figure S3**. Graph depict cholesterol levels in the medium of WT and *Mecp2* KO acutely isolated cortical astrocytes analysed through Amplex Red Assay. Statistical analysis was performed by Mann-Whitney test; n=6/9 exp.groups.

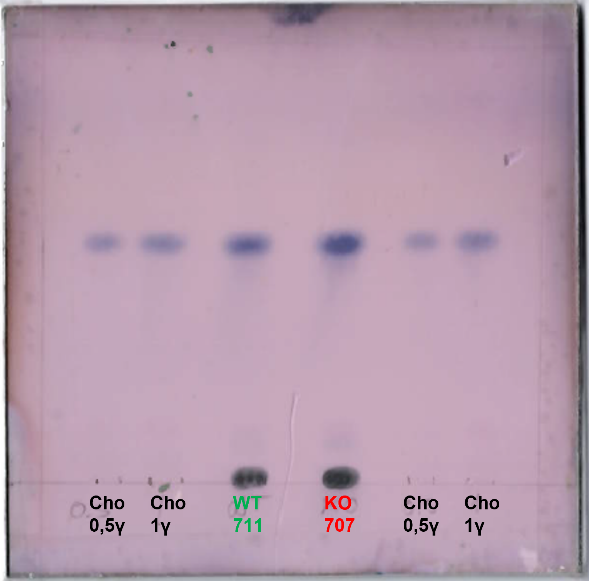

**Figure S4.** Representative HPTLC filter showing cholesterol signal in WT and *Mecp2* KO cerebral tissue. As control, known quantities of cholesterol were loaded in the same experiment.

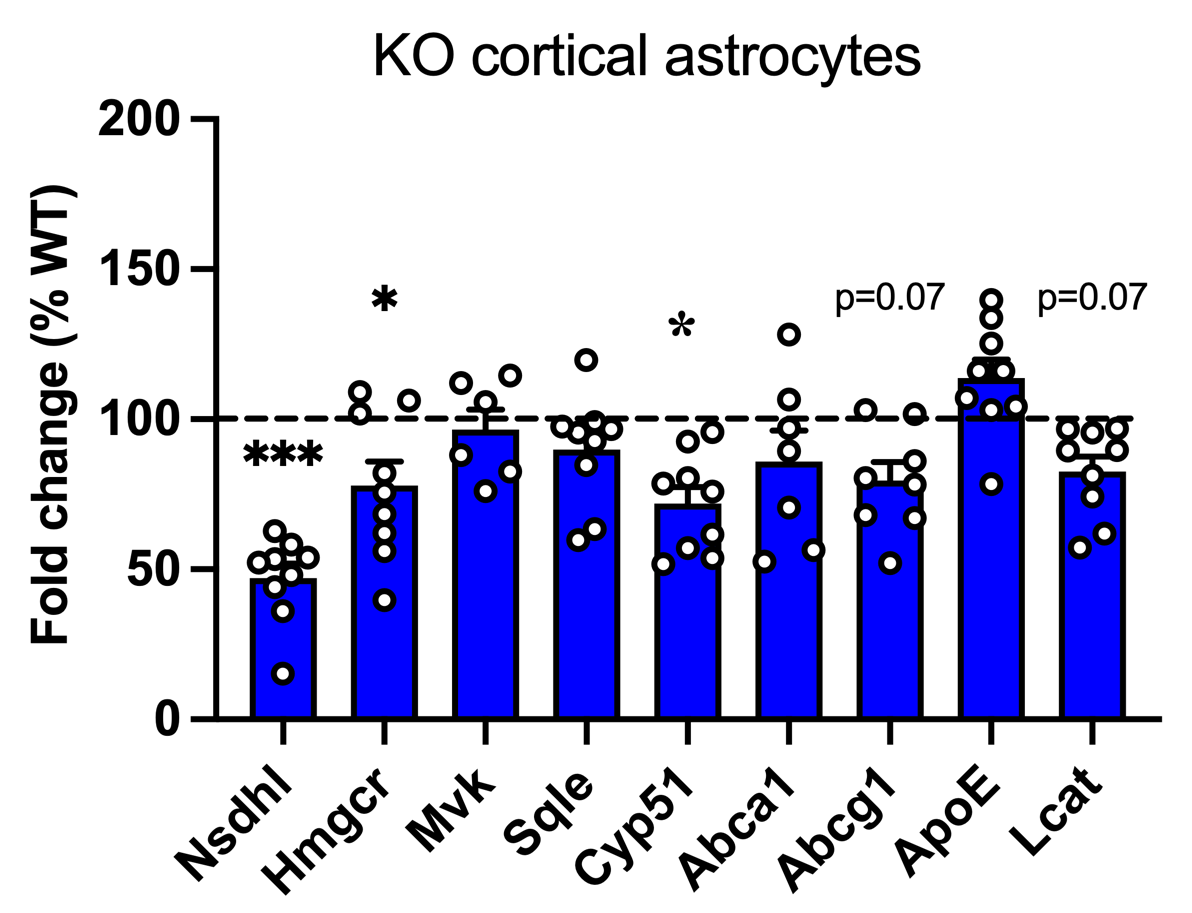

**Figure S5.** Histogram shows the mRNA levels of genes involved in the synthesis and transport of cholesterol in astrocytes isolated through MACS technology from the cortex of P40 mice. Data are expressed as percentages with respect to WT astrocytes, set at 100% and represented by the dotted line. Data are shown as mean±SEM. n=6/9 exp.groups. *p<0.05, ***p<0.001 by Student’s t-test.

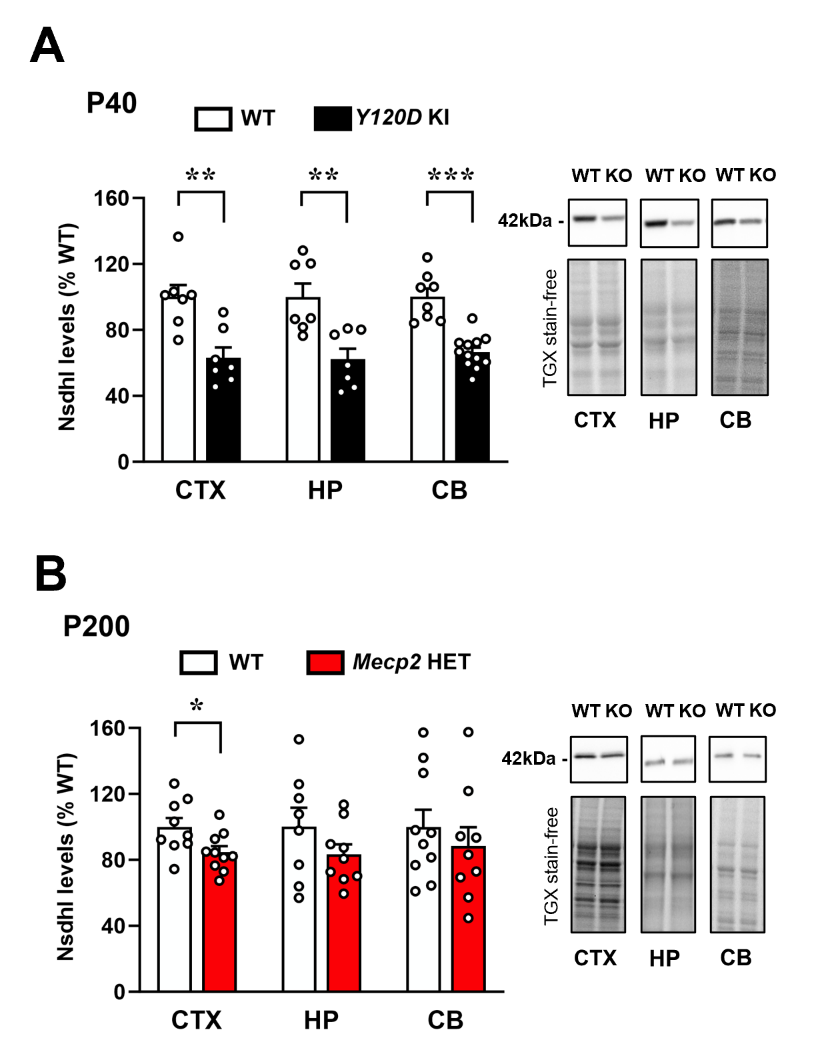

**Figure S6.** Histograms show the protein expression levels of Nsdhl measured in the cortex (CTX), hippocampus (HP) and cerebellum (CB) of KI Mecp2^Y120D/y^ (A) and *Mecp2* HET (B) animals. Data are expressed as percentages of corresponding WT animals and shown as mean±SEM. n= 8-10/exp.groups.  *p<0.05, **p<0.01, ***p<0.001 by Student’s t test. On the right of each graph, representative images of Nsdhl protein and total protein content visualized by TGX stain-free technology are shown.
